# Worldwide recombination in emergent white-nose syndrome pathogen *Pseudogymnoascus destructans*

**DOI:** 10.1101/868331

**Authors:** Lav Sharma, Márcia Sousa, Ana S. Faria, Margarida Nunes-Pereira, João A. Cabral, Alan J. L. Phillips, Guilhermina Marques, Maria das Neves Paiva-Cardoso

## Abstract

*Pseudogymnoascus destructans* (Pd), the emergent fungus causing bat “White-Nose Syndrome”, responsible for ∼6 million mortalities in the United States (US), is thought to expand clonally in North America and Europe. Presence of distinct mating-types in Europe led to numerous research attempts searching for population sexuality worldwide. This study not only presents the first evidence of genetic recombination in Pd but also detects recombination in Pd genotype data generated by previous studies in Europe and North America, through clone-corrected linkage disequilibrium analysis. Portuguese and other European populations are apparently reproducing through sex between two mating-types. Seeming parasexual recombination in the invasive single mating-type US population rings alarms for the North American bat populations and deserves urgent attention. This study emphasizes on clone-correction in linkage disequilibrium analysis.

**One Sentence Summary:** Clone-correction yielded signs of elusive recombination in the global “clonal” populations of white-nose syndrome pathogen.

## Main Text

White-nose syndrome (WNS), also called white-nose disease (WND) [1, 2, 3], is an emergent bat disease first reported in 2006 [4], caused by a novel fungal pathogen, *Pseudogymnoascus* (=*Geomyces*) *destructans* (Pd) [4, 5, 6]. *Pseudogymnoascus destructans* is a psychrophilic dermatophyte which has caused severe pathogenic effects in North American hibernating bats, leading up to catastrophic mortality rates, with over 6 million dead [3, 6, 7]. WND has turned into one of the most devastating wildlife epidemics in the recorded history of United States of America (US). Clinical signs of WND in bats include the distinctive presence of white fungal proliferation on glabrous skin of wings, ears and muzzle, cutaneous lesions, frequent arousal from torpor during hibernation, premature depletion of fat reserves, dehydration and electrolyte imbalance [1, 2, 4]. WND exhibits some of the worst possible epidemiological characteristics: (a) a highly virulent pathogen (Pd); (b) long-lasting persistence in hibernacula; (c) viability of conidia at elevated temperatures (24-37°C) suggesting survival on or within the bodies of bats and enabling long-distance dispersal; (d) a frequency and density-dependent transmission; and (e) a wide range of bat host species [8, 9].

In North America, WND has spread rapidly [10], having been identified in 33 US states and 7 Canadian provinces so far; meanwhile, Pd has been detected in 5 additional states. Bats represent approximately one-fifth of the total mammalian diversity [11], and as "key species’ in several ecosystems, play pivotal roles in plant pollination and control of insect populations, preventing huge losses in crops and forest production [12]. Over half of the 47 bat species living in the US and Canada depend on hibernation for winter survival [7]. Presently, 12 bat species, two of them endangered (*Myotis grisescens* and *Myotis sodalis*) and one threatened (*Myotis septentrionalis*), have been confirmed with WND in North America, which poses a risk for regional extinction in these populations [7, 13]. Hence, the massive bat population decline resulted from WND has caused severe ecological disequilibrium in ecosystems, and the loss of bats in North America could lead to agricultural losses of over $3.7 billion/year [13].

Globally, the fungus has been recorded in the majority of European countries and in China [1, 3, 8, 11, 14, 15], despite no mass mortality having been detected. In the Palearctic region, Pd has been detected in at least 21 bat species, highlighting its worldwide presence [7, 8, 15]. The majority of research studies corroborate the theory that Pd was introduced to North America from Europe [16, 17, 18], probably due to global trading or tourism. Its astonishingly rapid dispersion and epidemic character illustrates a classic case of the introduction of a highly lethal novel pathogen into a naïve host population, for example the well-known cases of *Batrachochytrium dendrobatidis* in amphibians, *Ophidiomyces ophiodiicola* in snakes, and *Cryptococcus gattii* in mammals [18-21].

Given the worldwide dispersion of Pd, understanding population structure and genetic diversity, and recombination is of utmost importance in understanding pathogen evolution, adaptation, and virulence [22, 23]. The presence of two different mating-types in same hibernacula in Europe, mating-type 1 (*MAT 1-1*) and mating-type 2 (*MAT 1-2*), hinted that Pd populations are undergoing meiosis [24]. However, numerous research attempts have been carried out worldwide and it has been noticed that the population expansion is clonal throughout. Phylogeographic studies based on strains collected from Eastern Europe and adjacent Russia [8], and from different Central and Western European countries across different years [17], did not report any recombination. Similarly, a clonal expansion of Pd has been reported repeatedly among different US states and Canadian provinces [12, 18, 25-28].

In the present study, a multilocus sequence typing (MLST) approach was undertaken using five loci to study the population genetics of 17 Pd isolates collected from hibernating bats in Portugal (Table S1). A total of eight haplotypes were found in the data (Fig. S1). Mean allelic diversity within Portuguese strains of Pd was 0.6176 ± 0.0556. Genetic diversity and related details, and phylogenetic trees based on each locus are presented in supplementary information (Table S2 and Figs. S2-S7). To identify evidences of genetic recombination among fungal populations, different tests were used. A concatenated alignment from five loci was analyzed using DnaSP version 5.10.01 [29] and at least one event of genetic recombination in history (*Rm* [30]) was detected among *MAT1-1* strains. Clear phylogenetic conflicts were noticed among different gene trees (Fig. 1), thus rejecting linkage disequilibrium.

**Figure1.**
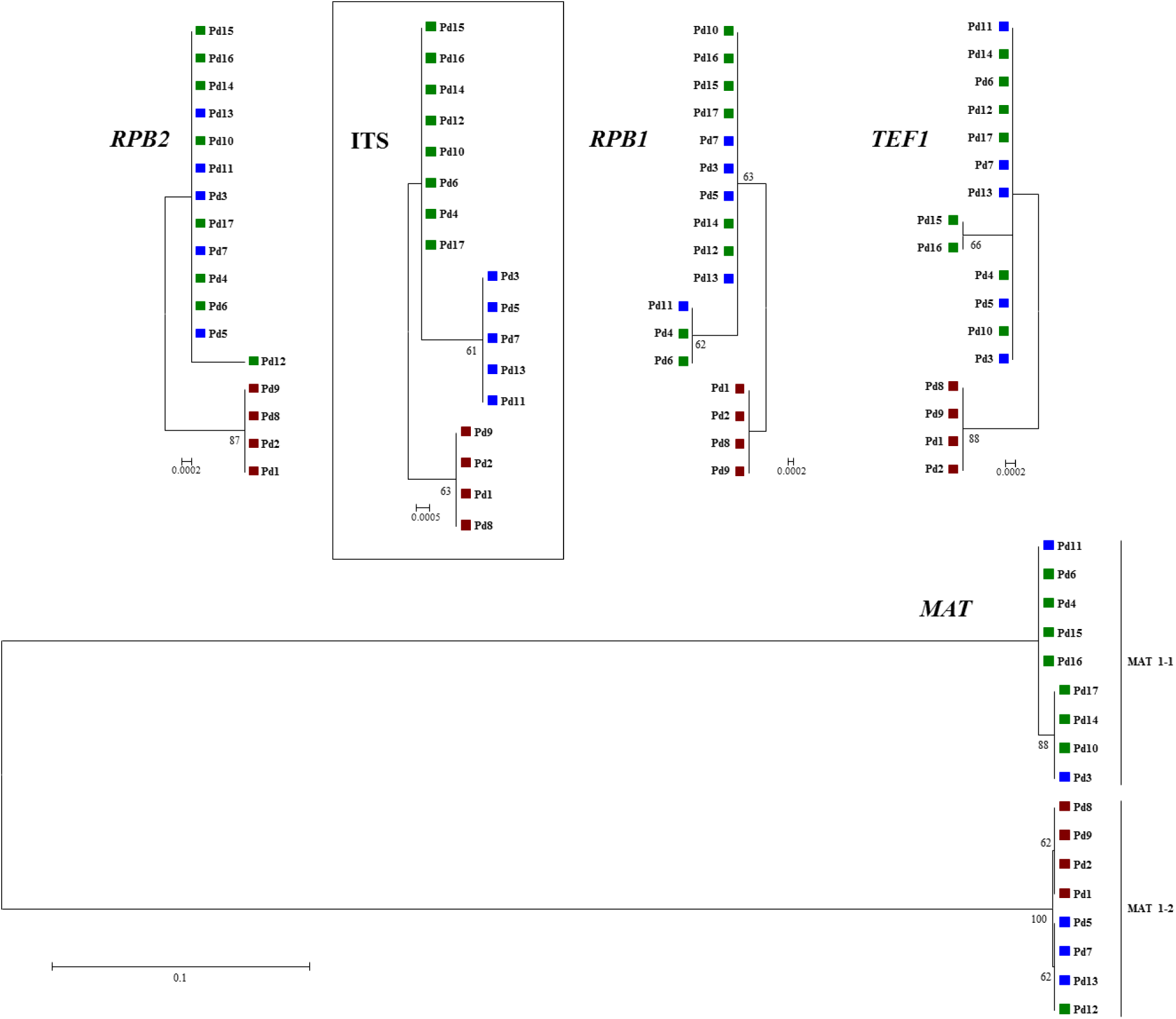
Phylogenetic incongruences among five loci trees. Unrooted Maximum-likelihood trees using Mega version 6.06 [33] were made using the best nucleotide substitution model, calculated with the in-built program. Nuclear ribosomal internal transcribed spacer (ITS) tree was used as the standard for the representation of the three different alleles, identified by different colors (green, blue and red).

Moreover, index of association (*I*_*A*_) and its modified statistics (*rBarD*) [31] were calculated to verify the null hypothesis of no linkage disequilibrium among loci, or recombination among the population from Portugal. Population haplotype data was analyzed with and without clone-correction as an earlier study has stressed on the sampling bias during fungal epidemics [23], and clone-correction has been successfully used in detecting recombination among populations of human and animal pathogenic fungi [19, 21]. In the five-locus based haplotype analysis of Portuguese Pd population, different sets with a minimum of three of more loci were tested for linkage disequilibrium analyses. No clone-correction failed to accept the null hypothesis of linkage equilibrium among loci (Tables 1 and S3). However, recombination was detected in the majority of the locus combinations while maintaining eight unique haplotypes in the clone-corrected data (Tables 1 and S4). Therefore, the clone-correction approach was further tested with the allele data from studies conducted earlier in different parts of Europe [8, 17]. Single nucleotide polymorphisms (SNPs) data from North America [28] was also tested.

**Table1:**
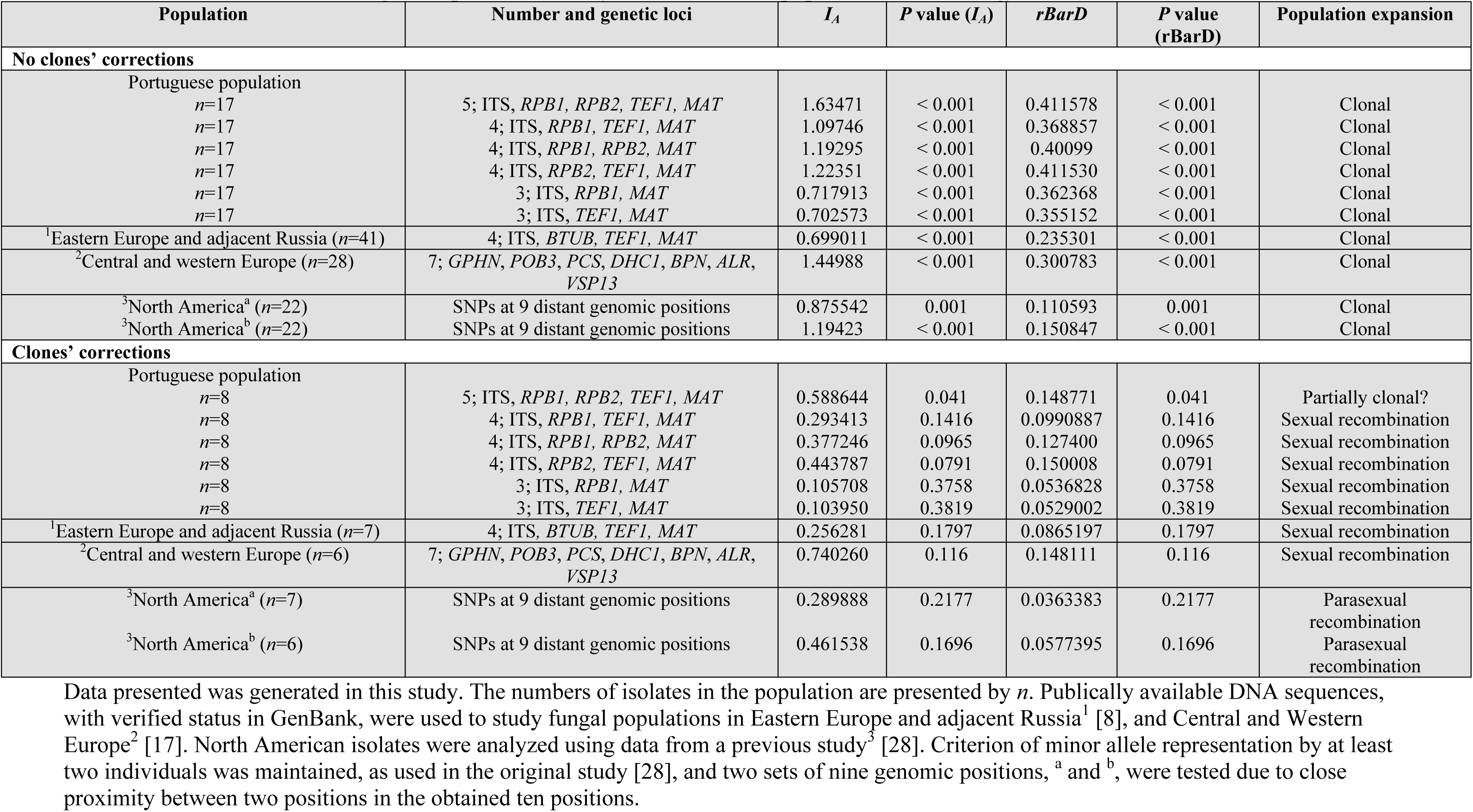
Linkage disequilibrium analyses of worldwide populations of *Pseudogymnoascus destructans*.

| Population | Number and genetic loci | $I_A$ | $P$ value ($I_A$) | $rBarD$ | $P$ value ($rBarD$) | Population expansion |
| --- | --- | --- | --- | --- | --- | --- |
| **No clones' corrections** |  |  |  |  |  |  |
| Portuguese population<br>$n=17$ | 5; ITS, *RPB1*, *RPB2*, *TEF1*, *MAT* | 1.63471 | < 0.001 | 0.411578 | < 0.001 | Clonal |
| $n=17$ | 4; ITS, *RPB1*, *TEF1*, *MAT* | 1.09746 | < 0.001 | 0.368857 | < 0.001 | Clonal |
| $n=17$ | 4; ITS, *RPB1*, *RPB2*, *MAT* | 1.19295 | < 0.001 | 0.40099 | < 0.001 | Clonal |
| $n=17$ | 4; ITS, *RPB2*, *TEF1*, *MAT* | 1.22351 | < 0.001 | 0.411530 | < 0.001 | Clonal |
| $n=17$ | 3; ITS, *RPB1*, *MAT* | 0.717913 | < 0.001 | 0.362368 | < 0.001 | Clonal |
| $n=17$ | 3; ITS, *TEF1*, *MAT* | 0.702573 | < 0.001 | 0.355152 | < 0.001 | Clonal |
| <sup>1</sup>Eastern Europe and adjacent Russia ($n=41$) | 4; ITS, *BTUB*, *TEF1*, *MAT* | 0.699011 | < 0.001 | 0.235301 | < 0.001 | Clonal |
| <sup>2</sup>Central and western Europe ($n=28$) | 7; *GPHN*, *POB3*, *PCS*, *DHC1*, *BPN*, *ALR*, *VSP13* | 1.44988 | < 0.001 | 0.300783 | < 0.001 | Clonal |
| <sup>3</sup>North America<sup>a</sup> ($n=22$) | SNPs at 9 distant genomic positions | 0.875542 | 0.001 | 0.110593 | 0.001 | Clonal |
| <sup>3</sup>North America<sup>b</sup> ($n=22$) | SNPs at 9 distant genomic positions | 1.19423 | < 0.001 | 0.150847 | < 0.001 | Clonal |
| **Clones' corrections** |  |  |  |  |  |  |
| Portuguese population<br>$n=8$ | 5; ITS, *RPB1*, *RPB2*, *TEF1*, *MAT* | 0.588644 | 0.041 | 0.148771 | 0.041 | Partially clonal? |
| $n=8$ | 4; ITS, *RPB1*, *TEF1*, *MAT* | 0.293413 | 0.1416 | 0.0990887 | 0.1416 | Sexual recombination |
| $n=8$ | 4; ITS, *RPB1*, *RPB2*, *MAT* | 0.377246 | 0.0965 | 0.127400 | 0.0965 | Sexual recombination |
| $n=8$ | 4; ITS, *RPB2*, *TEF1*, *MAT* | 0.443787 | 0.0791 | 0.150008 | 0.0791 | Sexual recombination |
| $n=8$ | 3; ITS, *RPB1*, *MAT* | 0.105708 | 0.3758 | 0.0536828 | 0.3758 | Sexual recombination |
| $n=8$ | 3; ITS, *TEF1*, *MAT* | 0.103950 | 0.3819 | 0.0529002 | 0.3819 | Sexual recombination |
| <sup>1</sup>Eastern Europe and adjacent Russia ($n=7$) | 4; ITS, *BTUB*, *TEF1*, *MAT* | 0.256281 | 0.1797 | 0.0865197 | 0.1797 | Sexual recombination |
| <sup>2</sup>Central and western Europe ($n=6$) | 7; *GPHN*, *POB3*, *PCS*, *DHC1*, *BPN*, *ALR*, *VSP13* | 0.740260 | 0.116 | 0.148111 | 0.116 | Sexual recombination |
| <sup>3</sup>North America<sup>a</sup> ($n=7$) | SNPs at 9 distant genomic positions | 0.289888 | 0.2177 | 0.0363383 | 0.2177 | Parasexual recombination |
| <sup>3</sup>North America<sup>b</sup> ($n=6$) | SNPs at 9 distant genomic positions | 0.461538 | 0.1696 | 0.0577395 | 0.1696 | Parasexual recombination |

Data presented was generated in this study. The numbers of isolates in the population are presented by $n$ . Publically available DNA sequences, with verified status in GenBank, were used to study fungal populations in Eastern Europe and adjacent Russia<sup>1</sup> [8], and Central and Western Europe<sup>2</sup> [17]. North American isolates were analyzed using data from a previous study<sup>3</sup> [28]. Criterion of minor allele representation by at least two individuals was maintained, as used in the original study [28], and two sets of nine genomic positions, <sup>a</sup> and <sup>b</sup>, were tested due to close proximity between two positions in the obtained ten positions.

By analyzing a previous European study [8], recombination was not found in the original data (Table S5), however, it was detected in clone-corrected haplotype data (Table S6) of *MAT1-1* and *MAT1-2* populations of Pd from Eastern Europe and adjacent Russia (*I*_*A*_=0.256281, *P*=0.1797; *rBarD* = 0.0865197, *P*=0.1797) (Table1). Similarly, while original data of Pd populations collected from several Western and Central European countries and along different years [17] (Table S7) failed to reject linkage disequilibrium, the clone-corrected multilocus data (Table S8) yielded signs of recombination (*I*_*A*_ = 0.74026, *P* = 0.116; *rBarD* = 0.148111, *P* = 0.116) (Table 1).

In North America, an earlier study reported clonal accumulation of mutations in 22 Pd strains [28]. To analyze recombination in the population history, they incorporated only those genomic positions where SNPs were reported in at least two strains, i.e., minor allele was represented at least two times (Table S9). We also considered this criterion. Moreover, to ensure accurate clone-correction, the genomic position with reoccurring missing information (14231511) was removed from the analyses. Out of the remaining ten positions, two of them, 4641354 and 4641566 were in very close proximity, and therefore two sets of nine positions were tested for linkage disequilibrium in North American Pd population, with and without clone-correction (Tables S9, S10 and S11). With all 22 strains, the clone-not-corrected data supported linkage disequilibrium among two sets of nine positions (*I*_*A*_ = 0.875542, *P* = 0.001; *rBarD* = 0.110593, *P* = 0.001) and (*I*_*A*_ = 1.19423, *P* < 0.001; *rBarD* = 0.150847, *P* < 0.001). However, clone-corrected data with seven strains and nine genomic positions (Table S10) rejected linkage disequilibrium among them (*I*_*A*_ = 0.289888, *P* = 0.2177; *rBarD* = 0.0363383, *P* = 0.2177). Similarly another set of clone-corrected data with six strains and nine positions (Table S11) resulted in the detection of genetic recombination in North-American populations (*I*_*A*_ = 0.461538, *P* = 0.1696; *rBarD* = 0.0577395, *P* = 0.1696) (Table 1).

Being the first study to report genetic evidence of sex in the causative agent of WNS or WND, since its discovery in 2006, highlights how little we know about the ecology of Pd. The present finding raises some relevant questions: (a) does the occurrence of sexual *MAT1-1* and *MAT1-2* recombination between strains of Pd throughout loom an outbreak in Europe?, (b) was WNS or WND also caused by a sudden invasion of a highly-virulent lineage that was recombining sexually in Europe, as was the case of the rapid expansion of frog chytridiomycosis through a hypervirulent recombining lineage of *Batrachochytrium dendrobatidis* [32]?

Nonetheless, this report rings an alarm bell for the remaining North American bat populations and requires urgent attention, as the only mating-type in North America has apparently started to recombine parasexually. A recent study has reviewed that the samplings during fungal epidemics are biased toward clones [23]. Clonal correction in linkage disequilibrium analysis among loci is a widely known methodology. Its application in neglecting linkage disequilibrium among different loci of otherwise primarily clonal Pd populations, worldwide, is another case of its usefulness in genotyping.

## Material and Methods

### Sample collections and isolation

Seventeen swabs were collected from infected or near-infected bats from three hibernacula located in Douro region of Portugal. Details about the location of the mines, and bat species and body part are provided TableS1. Collection procedure, fungal isolation, and DNA extraction protocols were adapted from previous studies [1, 34].

### Multilocus sequence typing approach

Multilocus sequence typing (MLST), a widely acknowledged genotyping methodology used to study populations of the human and animal fungal pathogens such as *Cryptococcus gattii, Batrachochytrium dendrobatidis, Metarhizium robertsii*, and *Pseudogymnoascus destructans* (Pd) [8, 17, 19, 21, 25-27, 35, 36]. The robustness of PCR-based MLST approach for *P. destructans* was earlier verified with respect to genome sequencing approaches, as the most common European haplotype based on eight loci [17], was an exact match to the respective loci with *Pseudogymnoascus destructans* (Pd) strains from North America [28]. Nonetheless, authors confirmed any substitution in this study multiple-time using different sets of PCR as suggested earlier [17]. Previous MLST studies based on eight loci, α-L-rhamnosidase (*ALR*); *Bpntase*, 3’(2’),5’-bisphosphate nucleotidase (*BPN*); Dynein heavy chain (*DHC1*); Gephyrin, molybdenum cofactor biosynthesis protein (*GPHN*); peroxisomal-coenzyme A synthetase (*PCS*); FACT complex subunit (*POB3*); vacuolar protein sorting-associated protein (*VPS13*); and signal recognition particle protein 72 (*SRP72*), reported no or limited resolution among isolates collected either in the US or among Central and Western European countries [17, 25-27]. A previous study used fungal taxonomic markers, such as nuclear ribosomal internal transcribed spacer (ITS), beta-tubulin (*BTUB*), translation elongation factor 1 alpha (*TEF1*); and mating type (*MAT*) locus (mating-type 1 (*MAT1-1*) or mating-type 2 (*MAT1-2*)) to study *P. destructans* populations in Eastern Europe and adjacent Russia [8]. Therefore, we also analyzed the regions of taxonomic markers, i.e., ITS, *TEF1*, largest subunit of the RNA polymerase II (*RPB1*), second largest subunit of the RNA polymerase II (*RPB2*); and, *MAT* locus (*MAT1-1* or *MAT 1-2*) to study Portuguese Pd populations.

### PCR amplification, DNA sequencing, and phylogenetic analyses

PCR amplification was performed using the MyTaq™ HS Mix (Bioline, Germany) following manufacturer’s guidelines. Details about the primers used to amplify the above-mentioned loci are provided in previous studies [8, 37]. PCR products were sequenced using the primers employed for their amplifications. Sequencing was performed by STABVIDA, Portugal. Sequences were trimmed and edited for their quality using BioEdit version 7.1.3.0 [38], and aligned using MAFFT version 7.222 [39], as described in [40]. Sequences were deposited in the European Nucleotide Archive (ENA) and the accession numbers are listed in Table S1. MEGA version 6.06 [33] was used to produce phylogenetic trees after choosing the right nucleotide substitution model using the in-built function. Branch support was calculated by 1000 bootstrap iterations. A local BLAST database was set up in Geneious version 9.1.6 (Biomatters Ltd., New Zealand) to retrieve sequences from publically available genome assembly of the Pd type strain 20631-21 (GCA_000184105) [41]; *Pseudogymnoascus pannorum* var. *pannorum* M1372 (GCA_000497305.1) [42]; *Pseudogymnoascus* sp. WSF 3629 (GCA_001662585.1), a strain from *P. roseus* sensu lato; and *Pseudogymnoascus* sp. 23342-1-I1 (GCA_001662575.1) [5, 41], for phylogeny. Sequences were also retrieved from GenBank and acknowledged subsequently.

### Linkage disequilibrium analysis, and recombination and neutrality tests

Index of association (*I*_*A*_) and the related statistics *rBarD* were calculated to test non-random association (multilocus linkage disequilibrium) in the digitalized alleles data from five loci originated in this study using MULTILOCUS version 1.3b [31]. To simulate complete panmixia, i.e., infinite recombination, 10,000 randomizations were performed. The null hypothesis of no linkage disequilibrium, or recombination, will not be rejected if the observed values of *I*_*A*_ and *rBarD* are not significantly different from the distribution of the values produced by 10,000 artificially recombining datasets [21, 31]. Allele data from this study and from previous studies in Europe and North America [8, 17, 28] were analyzed with and without clone-corrections, following its importance for successful use in detecting recombination among populations of animal pathogenic fungi in particular [19, 21, 36]. Subsequently, the values of *I*_*A*_ and *rBarD* were recalculated. An earlier study on recombination in insect fungal pathogen *M. robertsii* used three loci to detect recombination using *I*_*A*_ and *rBarD* statistics [36]. Therefore, a minimum set of three or more genetic loci from the Portuguese population were tested while maintaining eight unique haplotypes in the clone-corrected data. Digitalized allele data for the above mentioned studies are presented in the supplementary information (Tables S5-S11). DnaSp version 5.10.01 [29] was used to obtain information about individual loci; to infer minimum numbers of recombination events (*Rm* [30]) in the history of the populations; and to perform different neutrality tests, such as Tajima’s *D* [43], Fu & Li’s *F*\* [44], and Fu’s *F*_*S*_ [45] statistics, to estimate if there is any departure for any locus from the null hypothesis of neutral selection.

## ACKNOWLEDGMENTS

We would like to thank WNS research teams globally for working tirelessly while producing the data analyzed in this study. Present study would not be completed without their works. We also thank P. Barros and L. Braz (Laboratory of Applied Ecology, CITAB-UTAD) for help with bat handling, and I. Pinto and C. Teixeira for help with sample collection.

## Author contributions

Outlined the study: MdN.P-C., L.S., J.A.C., G.M.; collected samples and performed microbiological methods: M.S., A.S.F., M.N-P., MdN.P-C.; performed experiments: L.S., M.S., A.S.F., M.N-P.; analyzed and interpreted data: L.S., M.S.; wrote manuscript: L.S., MdN.P-C., A.S.F with assistance from A.J.L.P and G.M.

## Data and materials availability

All data are available in the main text or the supplementary materials. Requests for materials should be addressed to MdN.P-C.

