## Supplementary information for "Worldwide recombination in emergent white-nose syndrome pathogen *Pseudogymnoascus destructans*"

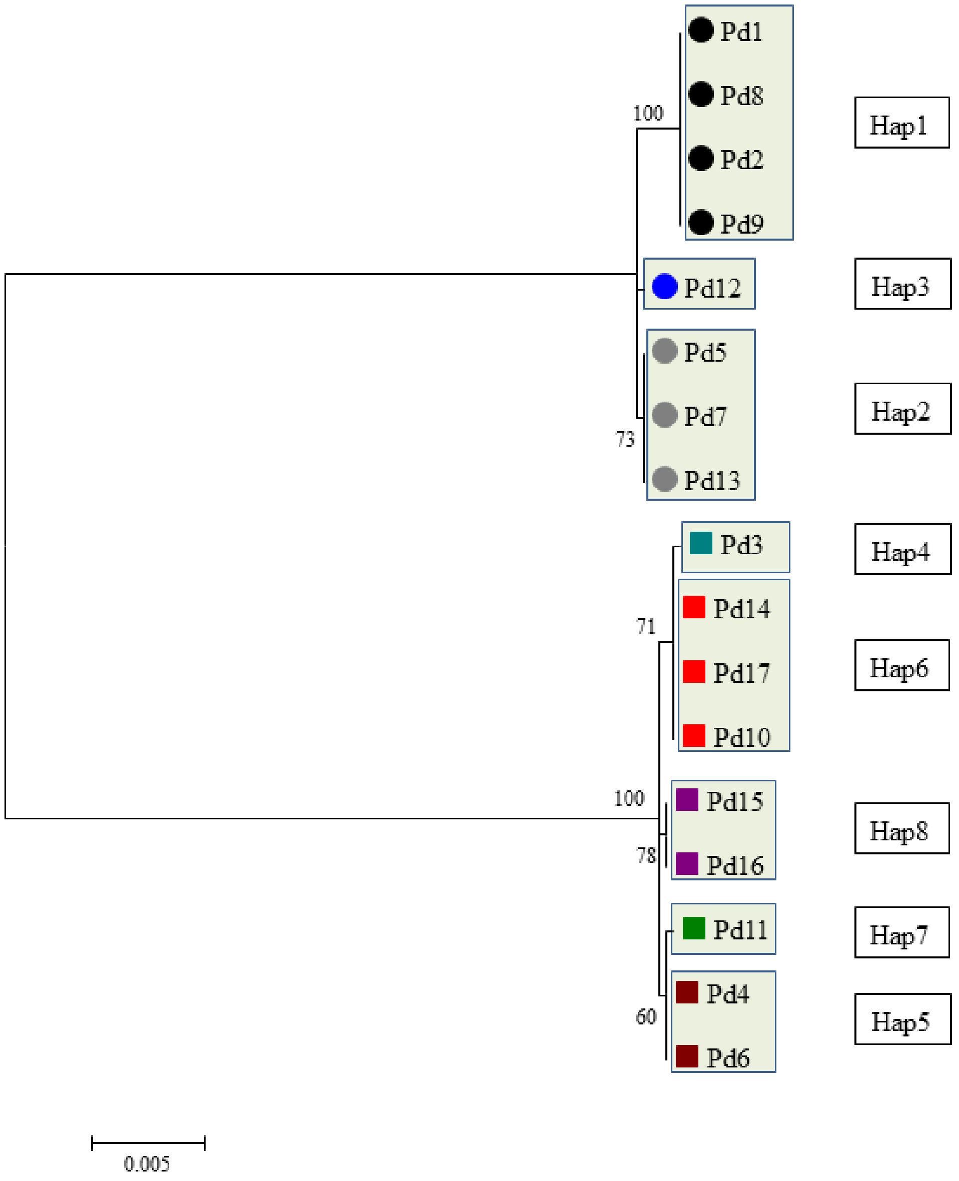


**Fig. S1. Unrooted Maximum-likelihood consensus tree of the 17 *Pseudogymnoascus destructans* isolates based on the concatenated DNA sequences of five MLST loci.** The tree was built with MEGA version 6.06 [33] using the Kimura 2-parameter model, considering all 3,732 positions in the alignment including gaps. Branch support values were obtained using 1,000 bootstrap replicates. The sequences presented with solid squares are *MAT1-1* isolates and solid circles are *MAT1-2* isolates, respectively. Each color and box represents a unique haplotype (Hap).

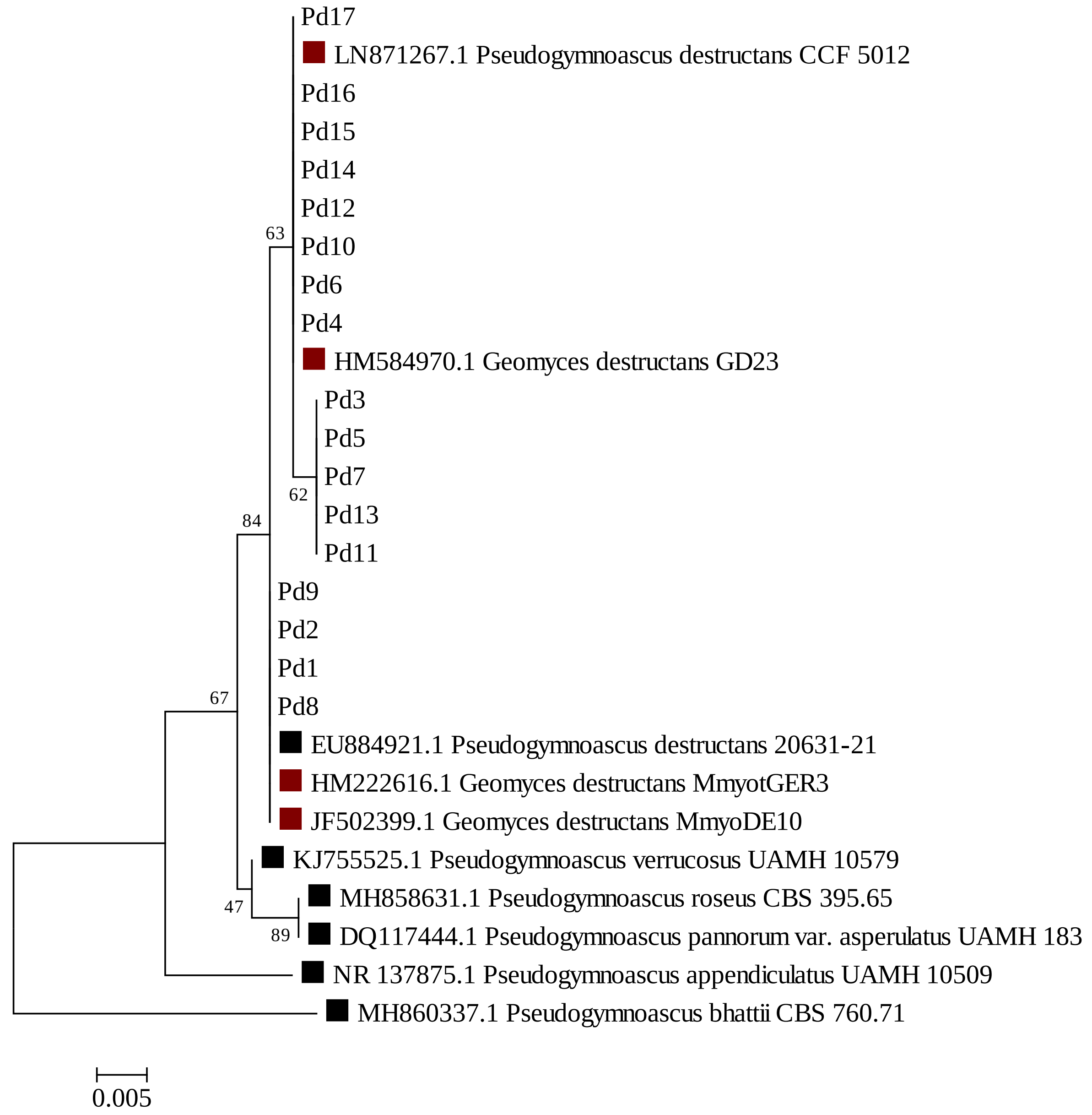


**Fig. S2. Maximum likelihood tree based on the DNA sequences of the nuclear ribosomal internal transcribed spacer (ITS).** The tree was built with MEGA version 6.06 [33] using the Kimura 2-parameter model, considering all 436 positions in the alignment. Branch support values were obtained using 1,000 bootstrap replicates. The sequences presented with solid squares were taken from public databases. Black squares represent type strains and crimson red squares represent other GenBank sequences.

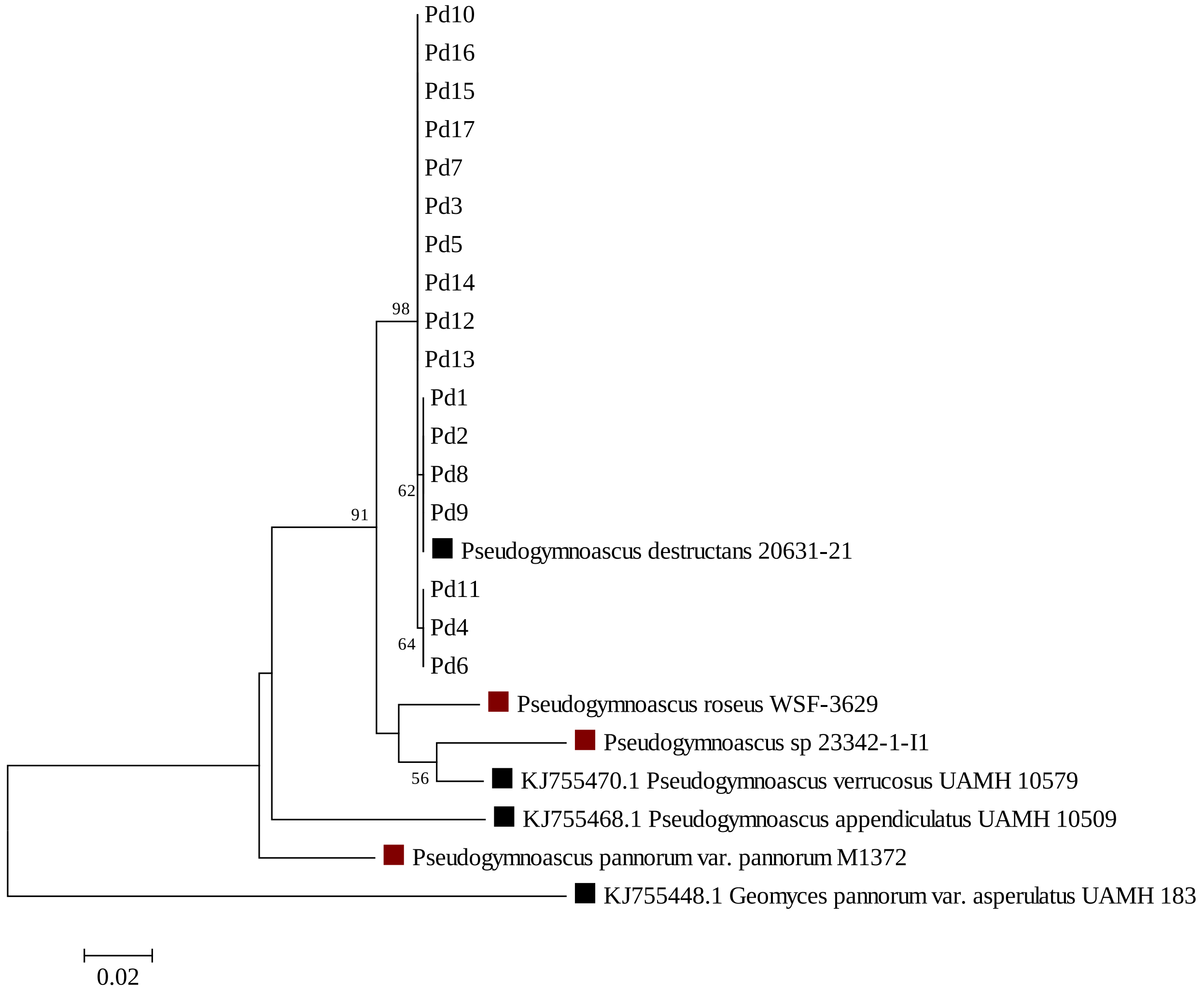


**Fig. S3. Maximum likelihood tree based on a region of the largest subunit of RNA polymerase II (*RPB1*) DNA sequences.** The tree was built with MEGA version 6.06 [33] using the Kimura 2-parameter model with a Gamma distribution, considering all 605 positions in the alignment. Branch support values were obtained using 1,000 bootstrap replicates. The sequences presented with solid squares were taken from public databases. Black squares represent type strains, and crimson red squares represent sequences retrieved from GenBank or publicly available genome assemblies.

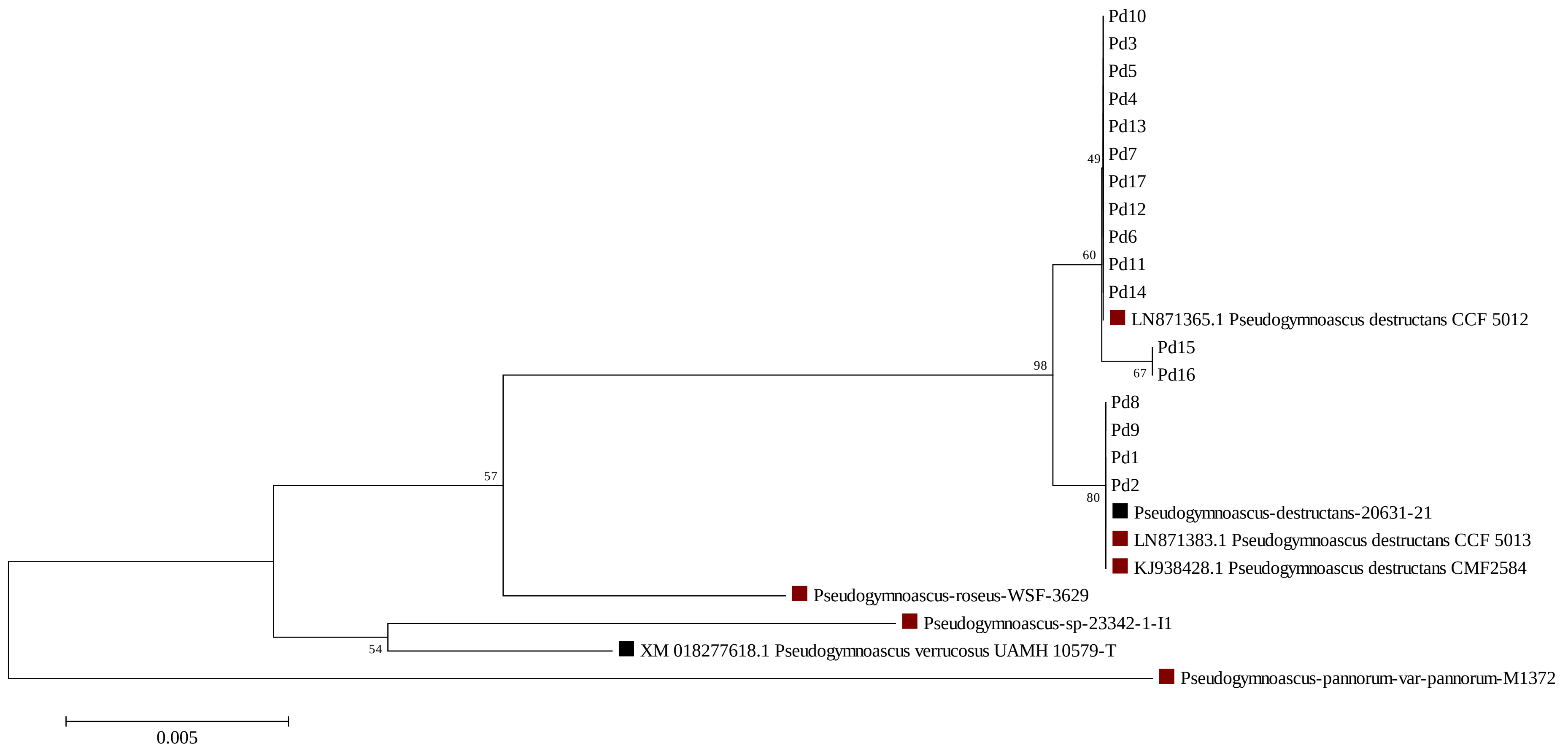


**Fig. S4. Maximum likelihood tree based on a region of the translation elongation factor 1 alpha (*TEF1*) DNA sequences.** The tree was built with MEGA version 6.06 [33] using the Tamura 3-parameter model with a Gamma distribution, considering all 912 positions in the alignment. Branch support values were obtained using 1,000 bootstrap replicates. The sequences presented with solid squares were taken from public databases. Black squares represent type strains, and crimson red squares represent sequences retrieved from GenBank or publicly available genome assemblies.

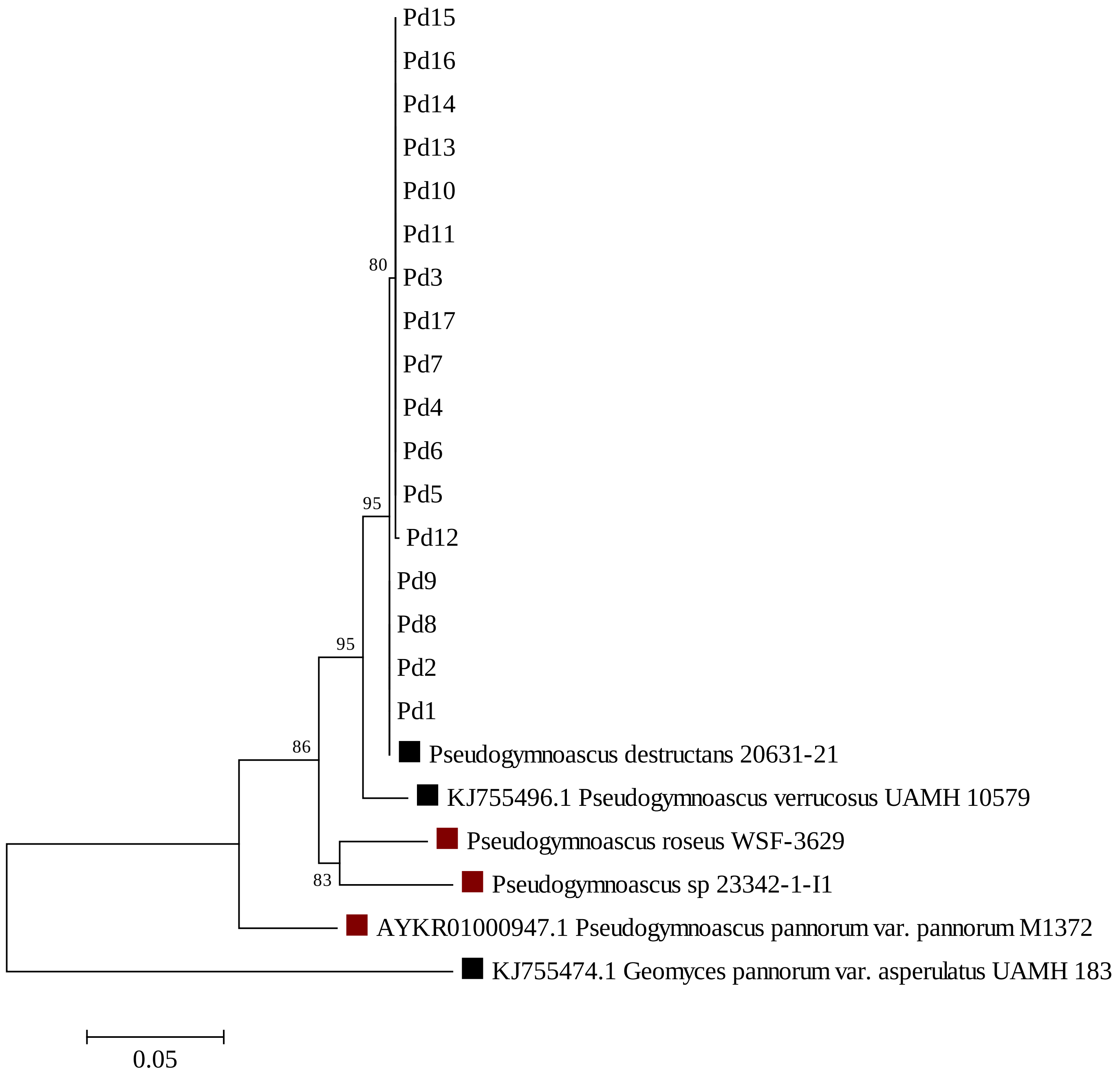


**Fig. S5. Maximum likelihood tree based on a region of the second largest subunit of RNA polymerase II (*RPB2*) DNA sequences.** The tree was built with MEGA version 6.06 [33] using the Kimura 2-parameter model with a Gamma distribution, considering all 912 positions in the alignment. Branch support values were obtained using 1,000 bootstrap replicates. The sequences presented with solid squares were taken from public databases. Black squares represent type strains, and crimson red squares represent sequences retrieved from GenBank or publicly available genome assemblies.

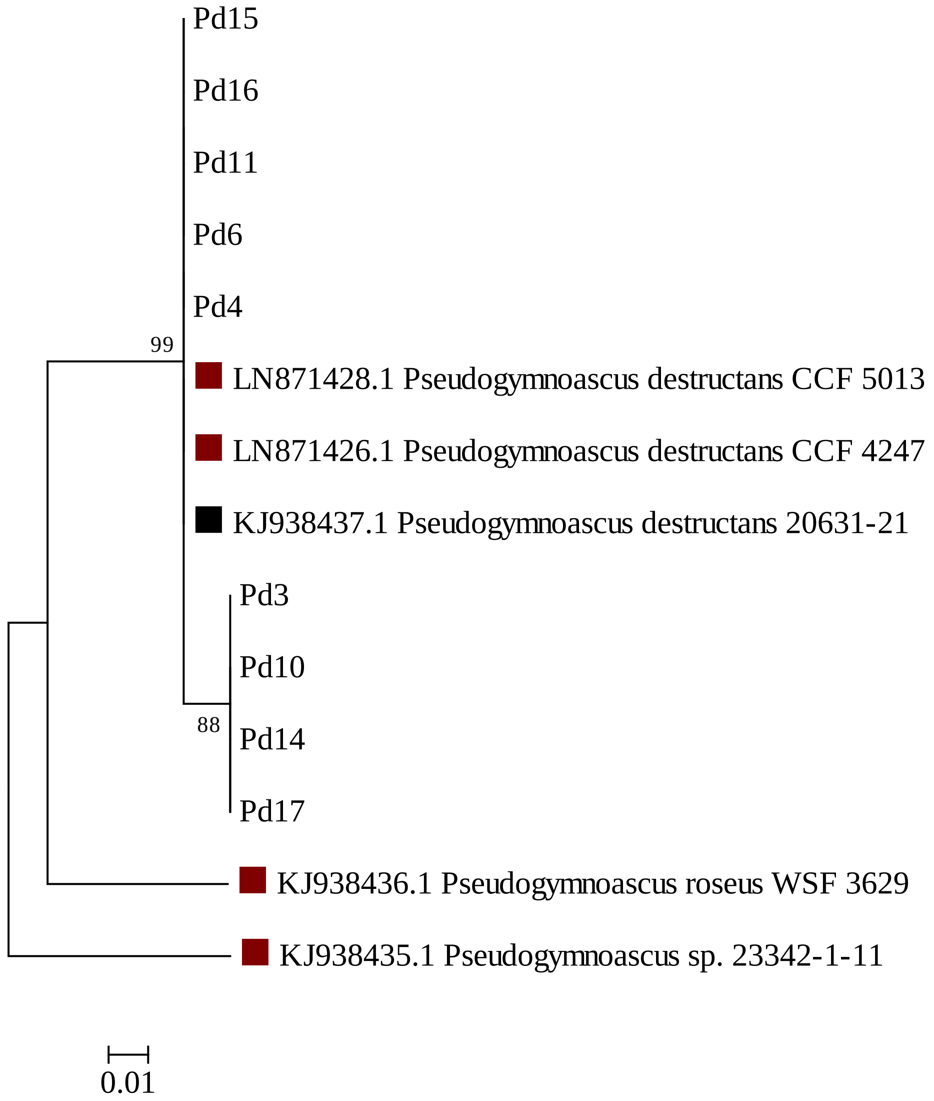


**Fig. S6. Maximum likelihood tree based on a region of the mating type 1 locus (*MAT-1-1*) DNA sequences.** The tree was built with MEGA version 6.06 [33] using the Jukes-Cantor parameter model with a Gamma distribution, considering all 182 positions in the alignment. Branch support values were obtained using 1,000 bootstrap replicates. The sequences presented with solid squares were taken from public databases. Black squares represent type strains, and crimson red squares represent sequences retrieved from GenBank or publicly available genome assemblies.

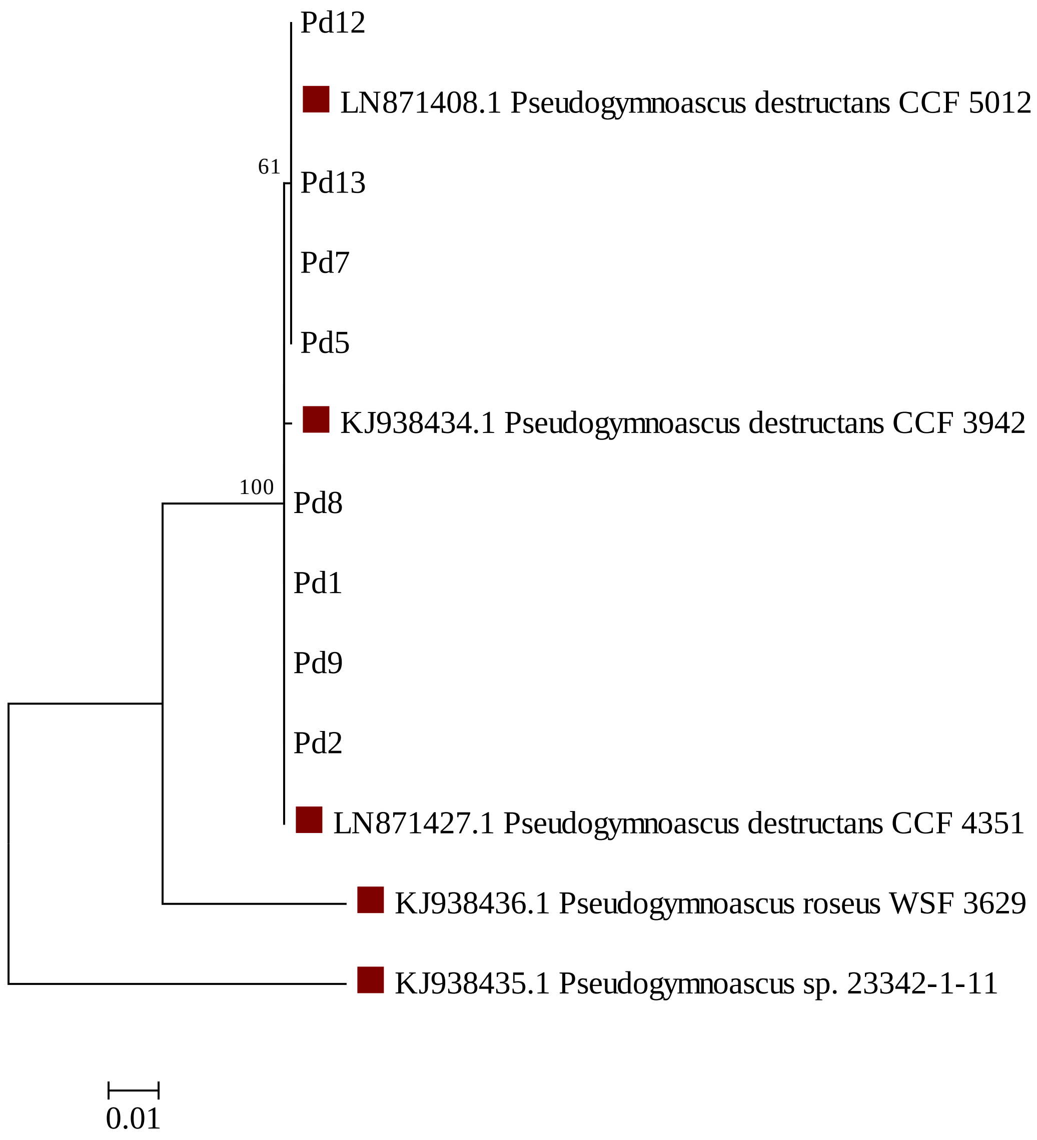


**Fig. S7. Maximum likelihood tree based on a region of the mating-type 2 locus (*MAT-1-2*) DNA sequences.** The tree was built with MEGA version 6.06 [33] using the Kimura 2-parameter model, considering all 709 positions in the alignment. Branch support values were obtained using 1,000 bootstrap replicates. The sequences presented with crimson red solid squares were taken from GenBank.

**Table S1. Strain information and generated accession numbers for each genetic locus.**

| Strains | Isolation source | Country | Area | Host | Mating Type | *MAT* | *TEF1* | *RPB1* | *RPB2* | ITS |
|---|---|---|---|---|---|---|---|---|---|---|
| Pd1 | Bat ear | Portugal | Mina Maria Isabel, Aldeia de Montes, Vila Real County, Vila Real District, Trás-os-Montes Region | *Myotis blythii* | *MAT 1-2* | LR736706 | LR736723 | LR736740 | LR736757 | LR739051 |
| Pd2 | Bat ear | Portugal | Mina Maria Isabel, Aldeia de Montes, Vila Real County, Vila Real District, Trás-os-Montes Region | *Myotis blythii* | *MAT 1-2* | LR736707 | LR736724 | LR736741 | LR736758 | LR739052 |
| Pd3 | Bat wing | Portugal | Mina Maria Isabel, Aldeia de Montes, Vila Real County, Vila Real District, Trás-os-Montes Region | *Myotis blythii* | *MAT 1-1* | LR736708 | LR736725 | LR736742 | LR736759 | LR739053 |
| Pd4 | Bat wing | Portugal | Mina Maria Isabel, Aldeia de Montes, Vila Real County, Vila Real District, Trás-os-Montes Region | *Myotis blythii* | *MAT 1-1* | LR736709 | LR736726 | LR736743 | LR736760 | LR739054 |
| Pd5 | Bat nose | Portugal | Mina Maria Isabel, Aldeia de Montes, Vila Real County, Vila Real District, Trás-os-Montes Region | *Myotis blythii* | *MAT 1-2* | LR736710 | LR736727 | LR736744 | LR736761 | LR739055 |
| Pd6 | Bat nose | Portugal | Mina Maria Isabel, Aldeia de Montes, Vila Real County, Vila Real District, Trás-os-Montes Region | *Myotis blythii* | *MAT 1-1* | LR736711 | LR736728 | LR736745 | LR736762 | LR739056 |
| Pd7 | Bat wing | Portugal | Mina Maria Isabel, Aldeia de Montes, Vila Real County, Vila Real District, Trás-os-Montes Region | *Myotis blythii* | *MAT 1-2* | LR736712 | LR736729 | LR736746 | LR736763 | LR739057 |
| Pd8 | Bat foot | Portugal | Mina Maria Isabel, Aldeia de Montes, Vila Real County, Vila Real District, Trás-os-Montes Region | *Myotis blythii* | *MAT 1-2* | LR736713 | LR736730 | LR736747 | LR736764 | LR739058 |
| Pd9 | Bat foot | Portugal | Mina Maria Isabel, Aldeia de Montes, Vila Real County, Vila Real District, Trás-os-Montes Region | *Myotis blythii* | *MAT 1-2* | LR736714 | LR736731 | LR736748 | LR736765 | LR739059 |
| Pd10 | Bat nose | Portugal | Minas de Campanhó, Campanhó Parish, Mondim de Basto County, Vila Real District, Trás-os-Montes Region | *Myotis blythii* | *MAT 1-1* | LR736715 | LR736732 | LR736749 | LR736766 | LR739060 |
| Pd11 | Bat wing | Portugal | Minas de Campanhó, Campanhó Parish, Mondim de Basto County, Vila Real District, Trás-os-Montes Region | *Myotis blythii* | *MAT 1-1* | LR736716 | LR736733 | LR736750 | LR736767 | LR739061 |
| Pd12 | Bat nose | Portugal | Minas de Campanhó, Campanhó Parish, Mondim de Basto County, Vila Real District, Trás-os-Montes | *Myotis blythii* | *MAT 1-2* | LR736717 | LR736734 | LR736751 | LR736768 | LR739062 |

| | | | | | | | | | | |
|---|---|---|---|---|---|---|---|---|---|---|
| | | | Region | | | | | | | |
| Pd13 | Bat wing | Portugal | Minas de Campanhó, Campanhó Parish, Mondim de Basto County, Vila Real District, Trás-os-Montes Region | *Myotis blythii* | *MAT 1-2* | LR736718 | LR736735 | LR736752 | LR736769 | LR739063 |
| Pd14 | Bat wing | Portugal | Minas de Campanhó, Campanhó Parish, Mondim de Basto County, Vila Real District, Trás-os-Montes Region | *Rhinolophus euryale* | *MAT 1-1* | LR736719 | LR736736 | LR736753 | LR736770 | LR739064 |
| Pd15 | Bat nose | Portugal | Minas de Campanhó, Campanhó Parish, Mondim de Basto County, Vila Real District, Trás-os-Montes Region | *Rhinolophus euryale* | *MAT 1-1* | LR736720 | LR736737 | LR736754 | LR736771 | LR739065 |
| Pd16 | Bat wing | Portugal | Minas de Campanhó, Campanhó Parish, Mondim de Basto County, Vila Real District, Trás-os-Montes Region | *Rhinolophus euryale* | *MAT 1-1* | LR736721 | LR736738 | LR736755 | LR736772 | LR739066 |
| Pd17 | Bat nose | Portugal | Túneis do Tua, Carrazeda de Ansiães County, Bragança District, Trás-os-Montes Region | *Miniopterus schreibersii* | *MAT 1-1* | LR736722 | LR736739 | LR736756 | LR736773 | LR739067 |

**Table S2. Loci information and neutrality tests as calculated by DnaSP version 5.10.01 [29].**

| **Genetic locus** | **Number of base pairs; gaps** | **Number of haplotypes (h)** | **Individual allelic diversity (hd)** | **Nucleotide diversity per site (pi)** | **Number of polymorphic sites** | **Number of parsimony informative sites** | **Tajima's *D*** | ***P* value (Tajima's D)** | **Fu & Li's *F**** | ***P value* (Fu & Li's *F**)** | **Fu's $F_S$** |
|---|---|---|---|---|---|---|---|---|---|---|---|
| ITS | 429; 0 | 3 | 0.676 ± 0.064 | 0.00192 ± 0.00029 | 2 | 2 | 0.9788 | > 0.1 | 1.04843 | > 0.1 | 0.638 |
| *TEF1* | 941; 0 | 3 | 0.544 ± 0.111 | 0.001 ± 0.00025 | 3 | 3 | 0.3125 | > 0.1 | 0.9628 | > 0.1 | 1.025 |
| *RPB1* | 652; 0 | 3 | 0.603 ± 0.098 | 0.001 ± 0.00022 | 2 | 2 | 0.4202 | > 0.1 | 0.8799 | > 0.1 | 0.283 |
| *RPB2* | 912; 0 | 3 | 0.471 ± 0.118 | 0.00097 ± 0.00026 | 3 | 2 | -0.0161 | > 0.1 | -0.5748 | > 0.1 | 0.7840 |
| *MAT1-1* | 325; 0 | 2 | 0.556 ± 0.090 | 0.00342 ± 0.00055 | 2 | 2 | 1.7542 | > 0.05 | 1.3550 | > 0.1 | 2.3020 |
| *MAT1-2* | 795; 0 | 2 | 0.571 ± 0.094 | 0.00072 ± 0.00012 | 1 | 1 | 1.444 | > 0.1 | 1.1003 | > 0.1 | 0.9660 |

**Table S3. Allele data generated using 17 *Pseudogymnoascus destructans* isolates collected from Portugal in this study.**

| | Locus → | ITS | *RPB1* | *RPB2* | *TEF1* | *MAT* | Haplotypes |
|---|---|---|---|---|---|---|---|
| **Strains** | | | | | | | |
| Pd1 | | 3 | 3 | 3 | 3 | 3 | Hap1 |
| Pd2 | | 3 | 3 | 3 | 3 | 3 | Hap1 |
| Pd3 | | 2 | 1 | 1 | 1 | 2 | Hap4 |
| Pd4 | | 1 | 2 | 1 | 1 | 1 | Hap5 |
| Pd5 | | 2 | 1 | 1 | 1 | 4 | Hap2 |
| Pd6 | | 1 | 2 | 1 | 1 | 1 | Hap5 |
| Pd7 | | 2 | 1 | 1 | 1 | 4 | Hap2 |
| Pd8 | | 3 | 3 | 3 | 3 | 3 | Hap1 |
| Pd9 | | 3 | 3 | 3 | 3 | 3 | Hap1 |
| Pd10 | | 1 | 1 | 1 | 1 | 2 | Hap6 |
| Pd11 | | 2 | 2 | 1 | 1 | 1 | Hap7 |
| Pd12 | | 1 | 1 | 2 | 1 | 4 | Hap3 |
| Pd13 | | 2 | 1 | 1 | 1 | 4 | Hap2 |
| Pd14 | | 1 | 1 | 1 | 1 | 2 | Hap6 |
| Pd15 | | 1 | 1 | 1 | 2 | 1 | Hap8 |
| Pd16 | | 1 | 1 | 1 | 2 | 1 | Hap8 |
| Pd17 | | 1 | 1 | 1 | 1 | 2 | Hap6 |

**Table S4. Clone-corrected allele data generated from *Pseudogymnoascus destructans* isolates collected from Portugal in this study.**

| | Locus → | ITS | *RPB1* | *RPB2* | *TEF1* | *MAT* | Haplotypes |
|---|---|---|---|---|---|---|---|
| **Strains** | | | | | | | |
| Pd1 | | 3 | 3 | 3 | 3 | 3 | Hap1 |
| Pd3 | | 2 | 1 | 1 | 1 | 2 | Hap4 |
| Pd4 | | 1 | 2 | 1 | 1 | 1 | Hap5 |
| Pd5 | | 2 | 1 | 1 | 1 | 4 | Hap2 |
| Pd10 | | 1 | 1 | 1 | 1 | 2 | Hap6 |
| Pd11 | | 2 | 2 | 1 | 1 | 1 | Hap7 |
| Pd12 | | 1 | 1 | 2 | 1 | 4 | Hap3 |
| Pd15 | | 1 | 1 | 1 | 2 | 1 | Hap8 |

**Table S5. Original allele data generated using a previous study on *Pseudogymnoascus destructans* isolates collected from Eastern Europe and adjacent Russia [8].**

| Locus → Strains† | *MAT* | *BTUB* | ITS | *TEF1* | Haplotype |
|---|---|---|---|---|---|
| CCF 3938 | 1 | 2 | 1 | 1 | Hap1 |
| CCF 4991 | 1 | 1 | 1 | 1 | Hap2 |
| CCF 4989 | 1 | 1 | 1 | 1 | Hap2 |
| CCF 5022 | 1 | 1 | 1 | 1 | Hap2 |
| CCF 5011 | 1 | 1 | 1 | 1 | Hap2 |
| CCF 4993 | 1 | 1 | 2 | 2 | Hap3 |
| CCF 5013 | 2 | 1 | 1 | 1 | Hap4 |
| CCF 4247 | 2 | 1 | 1 | 1 | Hap4 |
| CCF 5019 | 2 | 1 | 1 | 1 | Hap4 |
| CCF 4471 | 2 | 1 | 1 | 1 | Hap4 |
| CCF 5018 | 2 | 1 | 1 | 1 | Hap4 |
| CCF 5016 | 2 | 1 | 1 | 1 | Hap4 |
| CCF 5014 | 2 | 1 | 1 | 1 | Hap4 |
| CCF 4350 | 2 | 1 | 1 | 1 | Hap4 |
| AK-388-13 | 2 | 1 | 1 | 1 | Hap4 |
| CCF 5020 | 2 | 1 | 1 | 1 | Hap4 |
| CCF 4988 | 2 | 1 | 1 | 1 | Hap4 |
| CCF 5024 | 2 | 1 | 1 | 1 | Hap4 |
| CCF 5023 | 2 | 1 | 1 | 1 | Hap4 |
| CCF 4987 | 2 | 1 | 1 | 1 | Hap4 |
| CCF 4992 | 2 | 1 | 1 | 1 | Hap4 |
| CCF 5010 | 2 | 1 | 1 | 1 | Hap4 |
| CCF 4352 | 2 | 1 | 1 | 1 | Hap4 |
| CCF 4351 | 3 | 1 | 1 | 1 | Hap5 |
| CCF 4994 | 3 | 1 | 1 | 1 | Hap5 |
| CCF 3942 | 3 | 2 | 1 | 1 | Hap6 |
| CCF 5007 | 3 | 1 | 1 | 1 | Hap5 |
| CCF 5009 | 3 | 2 | 1 | 1 | Hap6 |
| CCF 4986 | 3 | 1 | 1 | 1 | Hap5 |
| CCF 5003 | 3 | 1 | 1 | 1 | Hap5 |
| CCF 4985 | 3 | 1 | 1 | 1 | Hap5 |
| CCF 3943 | 3 | 1 | 1 | 1 | Hap5 |
| CCF 5017 | 3 | 2 | 1 | 1 | Hap6 |
| CCF 4126 | 3 | 1 | 1 | 1 | Hap5 |
| CCF 5008 | 3 | 1 | 1 | 1 | Hap5 |
| CCF 4387 | 3 | 1 | 1 | 1 | Hap5 |
| CCF 5015 | 3 | 1 | 1 | 1 | Hap5 |
| CCF 5004 | 3 | 1 | 1 | 1 | Hap5 |
| CCF 5012 | 4 | 1 | 2 | 2 | Hap7 |
| CCF 5021 | 4 | 1 | 2 | 2 | Hap7 |
| AK-164-13 | 4 | 1 | 2 | 2 | Hap7 |

.†A total of 41 strains were found with unique data for all four loci.

**Table S6. Clone-corrected allele data generated using a previous study on *Pseudogymnoascus destructans* isolates collected from Eastern Europe and adjacent Russia [8].**

| Locus → Strains | *MAT* | *BTUB* | ITS | *TEF1* | Haplotype |
|---|---|---|---|---|---|
| CCF 3938 | 1 | 2 | 1 | 1 | Hap1 |
| CCF 4991 | 1 | 1 | 1 | 1 | Hap2 |
| CCF 4993 | 1 | 1 | 2 | 2 | Hap3 |
| CCF 5013 | 2 | 1 | 1 | 1 | Hap4 |
| CCF 4351 | 3 | 1 | 1 | 1 | Hap5 |
| CCF 3942 | 3 | 2 | 1 | 1 | Hap6 |
| CCF 5012 | 4 | 1 | 2 | 2 | Hap7 |

**Table S7. Original allele data generated using a previous study on 28 *Pseudogymnoascus destructans* isolates collected from Central and Western Europe along different years [17].**

| Locus† → Strains | *GPHN* | *POB3* | *PCS* | *DHC1* | *BPN* | *ALR* | *VSP13* | Haplotype |
|---|---|---|---|---|---|---|---|---|
| Gd_14 | 1 | 1 | 1 | 1 | 1 | 1 | 1 | Hap1 |
| Gd_16 | 1 | 1 | 1 | 1 | 1 | 1 | 1 | Hap1 |
| Gd_18 | 2 | 2 | 2 | 2 | 1 | 1 | 1 | Hap2 |
| Gd_21 | 1 | 1 | 1 | 1 | 1 | 1 | 1 | Hap1 |
| Gd_23 | 1 | 1 | 1 | 1 | 1 | 1 | 1 | Hap1 |
| Gd_24 | 1 | 1 | 1 | 1 | 1 | 1 | 1 | Hap1 |
| Gd_31 | 1 | 1 | 1 | 1 | 2 | 1 | 1 | Hap3 |
| Gd_35 | 1 | 1 | 1 | 1 | 1 | 1 | 1 | Hap1 |
| Gd_41 | 1 | 1 | 1 | 1 | 2 | 1 | 1 | Hap3 |
| Gd_44 | 1 | 1 | 1 | 1 | 2 | 1 | 1 | Hap3 |
| Gd_45a | 2 | 2 | 2 | 2 | 1 | 1 | 1 | Hap2 |
| Gd_46a | 1 | 1 | 1 | 1 | 1 | 1 | 1 | Hap1 |
| Gd_46b | 1 | 1 | 1 | 1 | 1 | 1 | 1 | Hap1 |
| Gd_46c | 1 | 1 | 1 | 1 | 1 | 1 | 1 | Hap1 |
| Gd_47a | 1 | 1 | 1 | 1 | 1 | 1 | 1 | Hap1 |
| Gd_48 | 1 | 1 | 1 | 1 | 1 | 2 | 1 | Hap4 |
| Gd_55 | 1 | 1 | 1 | 1 | 2 | 1 | 1 | Hap3 |
| Gd_62 | 1 | 1 | 1 | 1 | 3 | 1 | 1 | Hap5 |
| Gd_85 | 1 | 1 | 1 | 1 | 1 | 1 | 1 | Hap1 |
| Gd_94 | 1 | 1 | 1 | 1 | 1 | 1 | 1 | Hap1 |
| Gd_95 | 1 | 1 | 1 | 1 | 1 | 1 | 1 | Hap1 |
| Gd_97 | 1 | 1 | 1 | 1 | 1 | 1 | 1 | Hap1 |
| Gd_98 | 1 | 1 | 1 | 1 | 1 | 1 | 1 | Hap1 |
| Gd_99 | 1 | 1 | 1 | 1 | 1 | 1 | 1 | Hap1 |
| Gd_100 | 1 | 1 | 1 | 1 | 1 | 1 | 1 | Hap1 |
| Gd_101 | 1 | 1 | 1 | 1 | 1 | 1 | 1 | Hap1 |
| Gd_102 | 1 | 1 | 1 | 1 | 1 | 1 | 1 | Hap1 |
| Gd_105 | 1 | 1 | 1 | 3 | 1 | 1 | 1 | Hap6 |

†Data was analyzed using seven genes mentioned in the table. The gene *SRP72* was excluded as its GenBank status was unverified.

**Table S8. Clone-corrected allele data generated using a previous study on 28 *Pseudogymnoascus destructans* isolates collected from Central and Western Europe along different years [17].**

| Locus† → Strains | *GPHN* | *POB3* | *PCS* | *DHC1* | *BPN* | *ALR* | *VSP13* | Haplotype |
|---|---|---|---|---|---|---|---|---|
| Gd_14 | 1 | 1 | 1 | 1 | 1 | 1 | 1 | Hap1 |
| Gd_18 | 2 | 2 | 2 | 2 | 1 | 1 | 1 | Hap2 |
| Gd_44 | 1 | 1 | 1 | 1 | 2 | 1 | 1 | Hap3 |
| Gd_48 | 1 | 1 | 1 | 1 | 1 | 2 | 1 | Hap4 |
| Gd_62 | 1 | 1 | 1 | 1 | 3 | 1 | 1 | Hap5 |
| Gd_105 | 1 | 1 | 1 | 3 | 1 | 1 | 1 | Hap6 |

†Data was analyzed using seven genes mentioned in the table. The gene *SRP72* was excluded as its GenBank status was unverified.

**Table S9. Original single nucleotide polymorphism (SNP) data from 22 North American *Pseudogymnoascus destructans* strains with minor allele represented by at least two strains [28].**

| **Genomic positions →** | | | | | | | | | | | | **Haplotypes** |
|---|---|---|---|---|---|---|---|---|---|---|---|---|
| **Strains** | 4641354* | 4641566* | 8308033 | 11737596 | 12110873 | 13458202 | 23719871 | 24201679 | 25587530 | 5003057 | 14231511† | |
| 102203 | 0 | 0 | 0 | 0 | 0 | 0 | 1 | 0 | 1 | 0 | 0 | Hap1 |
| 192204 | 0 | 0 | 0 | 1 | 0 | 0 | 1 | 0 | 1 | 0 | 0 | Hap2 |
| 52201 | 0 | 0 | 0 | 0 | 0 | 0 | 1 | 0 | 1 | 0 | 0 | Hap1 |
| 671202 | 0 | 0 | 0 | 0 | 0 | 0 | 1 | 0 | 1 | 0 | 0 | Hap1 |
| 681103 | 0 | 0 | 0 | 0 | 0 | 0 | 1 | 0 | 1 | 0 | 0 | Hap1 |
| 692102 | 0 | 0 | 0 | 1 | 0 | 0 | 1 | 0 | 1 | 0 | 0 | Hap2 |
| 712206 | 0 | 0 | 0 | 1 | 0 | 0 | 1 | 0 | 1 | 0 | 1 | Hap3 |
| H07218 | 0 | 0 | 0 | 1 | 0 | 0 | 1 | 0 | 1 | 0 | 0 | Hap2 |
| M53205 | 0 | 0 | 0 | 0 | 0 | 0 | 1 | 0 | 1 | 0 | 0 | Hap1 |
| N_american | 0 | 1 | 0 | 0 | 0 | 0 | 0 | 0 | 0 | 0 | 0 | Hap4 |
| Nova_Scotia_1 | 1 | 0 | 0 | 1 | 0 | 0 | 1 | 0 | 1 | 0 | 1 | Hap5 |
| Nova_Scotia_2 | 1 | 0 | 0 | 1 | 0 | 0 | 1 | 0 | 1 | 0 | 1 | Hap5 |
| 20631.21 | 0 | 1 | 0 | 0 | 0 | 0 | 0 | 0 | 0 | 0 | 0 | Hap4 |
| 20631.008 | 0 | 0 | 0 | 0 | 0 | 0 | 1 | 0 | 1 | 1 | NC | ? |
| 26994.002 | 0 | 0 | 0 | 0 | 0 | 0 | 1 | 0 | 1 | 0 | NC | ? |
| 44797.145 | 0 | 0 | 0 | 0 | 0 | 0 | 1 | 0 | 1 | 0 | NC | ? |
| 27099.001 | 0 | 0 | 0 | 0 | 0 | 0 | 1 | 0 | 1 | 1 | NC | ? |
| UWMM_03 | 0 | 0 | 1 | 0 | 1 | 1 | 1 | 1 | 1 | 0 | 0 | Hap6 |
| UWMM_13 | 0 | 0 | 1 | 0 | 1 | 1 | 1 | 1 | 1 | 0 | 0 | Hap6 |
| UWMM_14 | 0 | 0 | 1 | 0 | 1 | 0 | 1 | 1 | 1 | 0 | 0 | Hap7 |
| WO2109 | 0 | 0 | 0 | 0 | 0 | 0 | 1 | 0 | 1 | 0 | 0 | Hap1 |
| X4148_13 | 0 | 0 | 0 | 1 | 0 | 0 | 1 | 0 | 1 | 0 | 0 | Hap2 |

*Positions were in close proximity and only one genomic position at a time was considered for linkage disequilibrium analyses. Genomic positions without any coverage for some strains in the original data are denoted by NC, as in the original study. †For an accurate clone-correction, position with missing information was also not considered in further analyses.

**Table S10. Clone-corrected† single nucleotide polymorphism (SNP) data set 1 generated from the previous study on North American *Pseudogymnoascus destructans* strains when the minor allele was represented by at least two strains in the original data [28].**

| **Genomic positions →** | | | | | | | | | **Haplotypes** |
|---|---|---|---|---|---|---|---|---|---|
| **Strains** | 4641354* | 8308033 | 11737596 | 12110873 | 13458202 | 23719871 | 24201679 | 25587530 | 5003057 | |
| 102203 | 0 | 0 | 0 | 0 | 0 | 1 | 0 | 1 | 0 | Hap1 |
| 192204 | 0 | 0 | 1 | 0 | 0 | 1 | 0 | 1 | 0 | Hap2 |
| N_american | 0 | 0 | 0 | 0 | 0 | 0 | 0 | 0 | 0 | Hap3 |
| Nova_Scotia_1 | 1 | 0 | 1 | 0 | 0 | 1 | 0 | 1 | 0 | Hap4 |
| 20631.008 | 0 | 0 | 0 | 0 | 0 | 1 | 0 | 1 | 1 | Hap5 |
| UWMM_03 | 0 | 1 | 0 | 1 | 1 | 1 | 1 | 1 | 0 | Hap6 |
| UWMM_14 | 0 | 1 | 0 | 1 | 0 | 1 | 1 | 1 | 0 | Hap7 |

. *The position was in close proximity with another position (4641566), and only one genomic position at a time was considered for linkage disequilibrium analyses.
†Position with missing information (14231511) was not considered for an accurate clone-correction.

**Table S11. Clone-corrected† single nucleotide polymorphism (SNP) data set 2 generated from the previous study on North American *Pseudogymnoascus destructans* strains when the minor allele was represented by at least two strains in the original data [28].**

| **Genomic positions →** | | | | | | | | | **Haplotypes** |
|---|---|---|---|---|---|---|---|---|---|
| **Strains** | 4641566* | 8308033 | 11737596 | 12110873 | 13458202 | 23719871 | 24201679 | 25587530 | 5003057 | |
| 102203 | 0 | 0 | 0 | 0 | 0 | 1 | 0 | 1 | 0 | Hap1 |
| 192204 | 0 | 0 | 1 | 0 | 0 | 1 | 0 | 1 | 0 | Hap2 |
| N_american | 1 | 0 | 0 | 0 | 0 | 0 | 0 | 0 | 0 | Hap3 |
| 20631.008 | 0 | 0 | 0 | 0 | 0 | 1 | 0 | 1 | 1 | Hap4 |
| UWMM_03 | 0 | 1 | 0 | 1 | 1 | 1 | 1 | 1 | 0 | Hap5 |
| UWMM_14 | 0 | 1 | 0 | 1 | 0 | 1 | 1 | 1 | 0 | Hap6 |

*The position was in close proximity with another position (4641354), and only one genomic position at a time was considered for linkage disequilibrium analyses. †Position with missing information (14231511) was not considered for an accurate clone-correction.
